# On the logic of Fisherian sexual selection

**DOI:** 10.1101/815613

**Authors:** Carl Veller, Pavitra Muralidhar, David Haig

## Abstract

In Fisher’s model of sexual selection, a female preference for a male trait spreads together with the trait because their genetic bases become correlated. This can be interpreted as a ‘greenbeard’ system: a preference gene, by inducing a female to mate with a trait-bearing male, favors itself because the male is disproportionately likely also to carry the preference gene. Here, we use this logic to argue that Fisherian sexual selection in diploids proceeds via two channels, corresponding to two reasons that trait-bearing males disproportionately carry preference genes: (i) trait-bearing males are disproportionately the product of matings between preference-bearing mothers and trait-bearing fathers, and thus trait and preference genes are correlated ‘in trans’; (ii) trait and preference genes come into gametic phase disequilibrium, and thus are correlated ‘in cis’. Gametic phase disequilibrium is generated by three distinct mechanisms: a ‘recombination mechanism’, a ‘dominance mechanism’, and a ‘sexual admixture mechanism’. The trans channel does not operate when sexual selection is restricted to the haploid phase, and therefore represents a fundamental difference between haploid and diploid models of sexual selection. We use simulation experiments to artificially eliminate the cis channel, and show that a preference gene can spread in its absence in the diploid model, but not in the haploid model. We further show that the cis and trans channels contribute equally to the spread of the preference when recombination between the preference and trait loci is free, but that the trans channel becomes substantially more important when linkage is tight.

## Introduction

It is an evolutionary paradox that males often display costly ornaments or behaviors and females often prefer such males in mate choice. In *The Genetical Theory of Natural Selection*, Fisher (1930) outlined a theory for how such a state might evolve. In Fisher’s theory of sexual selection, (i) if the female mate preference becomes sufficiently common, the sexual advantage it confers on the male ornament/behavior (the male ‘trait’) outweighs the male trait’s survival cost, and (ii) the resulting positive selection of the male trait causes indirect positive selection of the female preference, as their genetic bases become correlated because of non-random mating. Thus, both trait and preference spread together.

Half a century later, the possibility of Fisherian sexual selection was demonstrated mathematically by Lande (1981), assuming a quantitative genetic basis of preference and trait, and by Kirkpatrick (1982), assuming a haploid population genetics model where preference and trait are each encoded at single loci. A major advantage of the population genetics approach is that the dynamics can be analyzed in full generality (Kuijper, Pen & Weissing 2012), whereas analysis of quantitative genetic models such as Lande’s typically requires simplifying assumptions, for example a pre-existing, constant genetic correlation between preference and trait.

Since we are usually interested in mate choice and sexual selection in diploid organisms, an important question is whether the population dynamics differs between the two-locus haploid model—studied by Kirkpatrick (1982) for its mathematical tractability—and the diploid analog. Previous work has uncovered differences, particularly in the structure of sets of equilibria possible under the two models (Heisler & Curtsinger 1990, Gomulkiewicz & Hastings 1990, Otto 1991, Greenspoon & Otto 2009). For example, stable paths of equilibria involving polymorphism at both loci are characteristic of the haploid model, but are not observed for most configurations of the diploid model (Heisler & Curtsinger 1990). In addition, intermediate dominances of the trait allele in its negative viability effect and its positive effect on attractiveness can combine to produce overall fitness overdominance (Curtsinger & Heisler 1988). This phenomenon, which is obviously not possible in the haploid model, promotes long-term polymorphism at the trait locus. In general, however, it has been argued that many of the differences between the diploid and haploid models are due to the greater complexity of the diploid model—both in the number of parameters and the number of possible genotypes—rather than sexual selection occurring in the diploid phase *per se* (Barton & Turelli 1991).

Here, we make use of the greenbeard interpretation of Fisherian sexual selection (Dawkins 1986; Pizzari & Gardner 2012; Faria, Varela & Gardner 2018) to compare the haploid and diploid models, and to gain a greater conceptual understanding of the general operation of Fisher’s process. A greenbeard gene (Dawkins 1976) causes its bearer both to grow a green beard and to behave altruistically towards others with green beards. The gene in a potential benefactor uses the green beard of potential beneficiaries as information that they also carry the gene; this allows the gene to favor copies of itself in other individuals, and thus to spread. In Fisherian sexual selection, a preference allele causes its female bearers to behave ‘altruistically’ towards males who display the trait—by favoring them in mating. If males displaying the trait are disproportionately likely to carry the preference allele, then, by conferring a mating advantage on such males, the preference allele confers a mating advantage on copies of itself. In this way, the trait acts as a green beard, providing information that its bearers likely carry the preference allele.

The greenbeard interpretation thus brings into sharp relief the key question underlying Fisher’s theory of sexual selection: Why are trait-displaying males disproportionately likely to carry the preference allele? That is, what is the nature of the association between the preference and trait alleles? When sexual selection occurs in the haploid phase, as in the model of Kirkpatrick (1982), the only possible association between the trait and preference alleles in males is one of gametic phase disequilibrium, or cis-linkage disequilibrium. However, if sexual selection occurs in the diploid phase, there are two possible associations between the trait and preference alleles: they can be in cis-linkage disequilibrium and/or in trans-linkage disequilibrium (Fig. 1). Trans-linkage disequilibrium is a direct consequence of the action of the mate preference, which causes a disproportionate number of offspring to inherit the preference allele from their mothers and the trait allele from their fathers. Cis-linkage disequilibrium is generated by a network of interacting mechanisms that we explore below.

**Figure 1.**
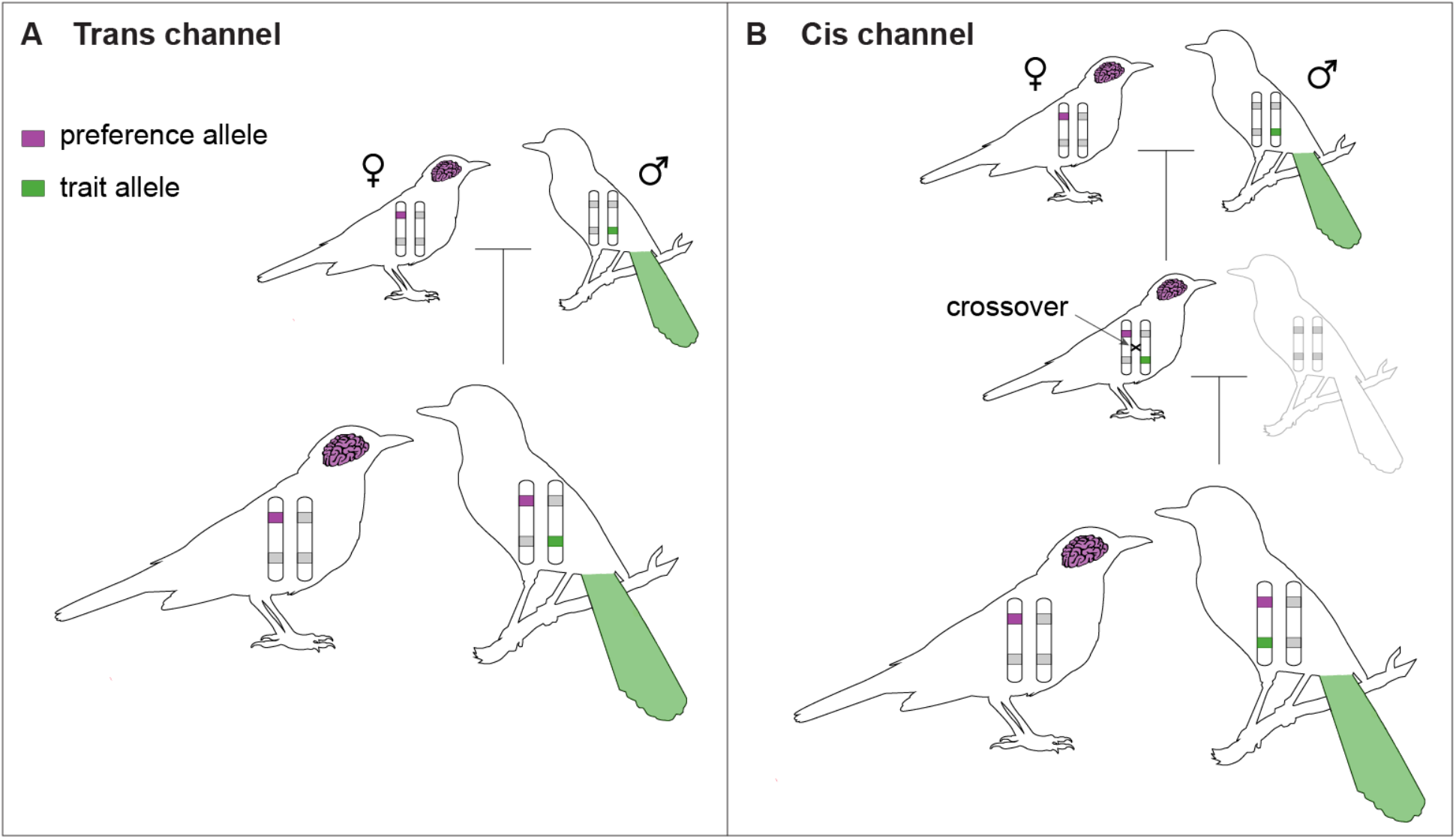
The trans and cis channels of Fisherian sexual selection in diploids. Under the greenbeard interpretation of Fisherian sexual selection, a preference allele acting in a female uses a male’s display of the preferred trait (here, a green tail) as information that he probably carries the preference allele. This requires a statistical association between the trait allele and the preference allele. Two such associations exist. **A**. Because a trait-displaying male is disproportionately likely to be the product of a mating between a preference-bearing mother and a trait-displaying father, he is disproportionately likely to carry the preference and trait alleles in a trans association. These trans associations are the basis for the trans channel of Fisherian sexual selection in diploids. **B**. By the action of the mate preference in the grandparental generation, individuals in the parental generation are disproportionately likely to carry the preference and trait alleles in a trans association. This gives the alleles disproportionate opportunities to recombine into the same gamete, and thus to be carried in a cis association in males in the current generation. Two other mechanisms, discussed in the text, also generate cis associations between the preference and trait alleles. These cis associations are the basis for the cis channel of Fisherian sexual selection in diploids. The two loci are displayed as being on the same chromosome here, but could also be on separate chromosomes.

These two genetic associations between the trait and preference represent two channels by which Fisherian sexual selection can operate in diploids. In this paper, we demonstrate the independent operation of the trans channel in diploid sexual selection, and investigate what factors influence the relative importance of the trans and cis channels to the spread of the preference. Our aim is to gain a conceptual understanding of the phenomena we discuss, and for this reason we first develop the logic of our arguments and then illustrate this logic with a few instructive simulations, rather than constructing the arguments on the basis of a simulation scan over the entire parameter space. Our primary focus is on the non-equilibrium dynamics of sexual selection, for several reasons: (i) the equilibria of the haploid and diploid models are well studied (Kirkpatrick 1982, Heisler & Curtsinger 1990, Gomulkiewicz & Hastings 1990), (ii) in many configurations of these models, the dynamics are sufficiently slow that non-equilibrium dynamics are important even on long timescales (e.g., Heisler & Curtsinger 1990, Otto & Greenspoon 2009, and below), and (iii) as we show, the non-equilibrium dynamics are often more revealing of the mechanisms underlying Fisherian sexual selection.

## The model

The model we employ is the diploid version of the canonical two-locus model of Fisherian sexual selection (Kirkpatrick 1982), as studied, for instance, by Gomulkiewicz & Hastings (1990), Heisler & Curtsinger (1990), Otto (1991), and Greenspoon & Otto (2009). The organism is sexual, and both natural and sexual selection are restricted to the diploid phase of its life cycle. Generations are non-overlapping, and the population is large enough that dynamics can be assumed to be deterministic.

There are two autosomal loci, the ‘trait locus’ and the ‘preference locus’, which recombine with rate *r*. Two alleles segregate at the trait locus: the wild-type *t* and the ‘trait allele’ *T*. *T* encodes a male-specific trait which reduces the viability of its bearers by a factor 1– *h*_*T*_*s*_*T*_ for *Tt* males and 1– *s*_*T*_ for *TT* males, where *h*_*T*_ is the dominance of *T* with respect to *t*. Two alleles segregate at the preference locus: the wild-type *p* and the ‘preference allele’ *P*. *P* encodes a female mate preference for the male trait, according to the usual ‘fixed relative preference’ model (Seger 1985). A *PP* female is more likely to mate with a given *TT* male or a given *Tt* male than with a given *tt* male by factors *α* and *α*^*hr*^ respectively, where *α* > 1 is the strength of the preference. The analogous factors for a *Pp* female are 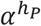 and 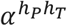 where *h*_*p*_ is the dominance of *P* with respect to *p*. For our main results, we assume co-dominance at both loci (*h*_*r*_ = *h*_*p*_ = 1/2), though we also examine the effects of different dominance values of *T*. For an explanation of the exponential functional relationship between preference strength and dominance coefficients, see Muralidhar (2019). Alleles at the preference locus have no direct influence on viability, fertility, or fecundity, although we briefly consider the effect of intrinsically costly preferences in the Discussion. The alleles at the trait locus mutate to each other with symmetric rate *μ* per replication.

Each generation, viability selection acts on juveniles, after which mating occurs subject to the preferences described above. This leads to a standard set of recursions describing the population dynamics (e.g., Greenspoon & Otto 2009). Our results derive from computer simulations of these recursions. Unless otherwise stated, we begin each simulation with the two loci in Hardy-Weinberg and linkage equilibrium.

The canonical two-locus haploid model of Fisherian sexual selection (Kirkpatrick 1982) is the same as that described above, except that natural and sexual selection are restricted to the haploid phase of the life cycle (which makes the dominance coefficients *h*_*T*_ and *h*_*p*_ irrelevant).

### Selection on the preference allele

Fig. 2A shows some typical frequency trajectories of the preference allele *P* and the trait allele *T* in the diploid model of Fisherian sexual selection. Several general features can be observed from these trajectories. First, *P* increases in frequency only when *T* also increases in frequency. This is because selection on *P* is indirect, being a result of selection on *T* and the fact that *P* and *T* become associated. Second, when the trait reduces the viability of its male bearers, the preference must initially be sufficiently common for *T* (and thus, indirectly, *P*) to be positively selected. Third, increases in frequency of *P* tend to be transient and small, as *T* is rapidly driven to fixation.

**Figure 2.**
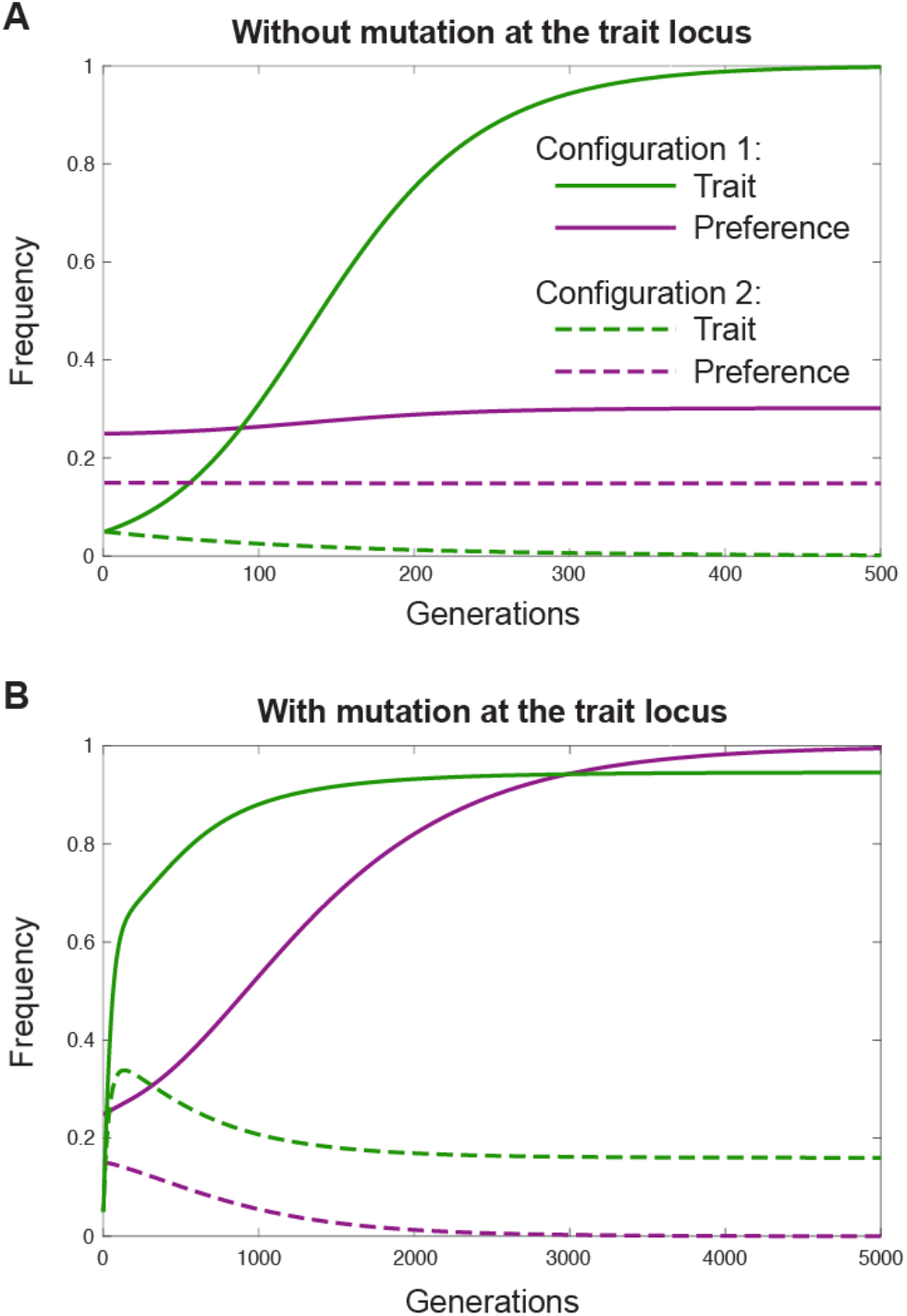
Mutation at the trait locus allows long-term spread of the preference. Frequency trajectories of the trait and preference alleles, for two initial starting frequencies of the preference allele. In configuration 1, the preference is initially common enough that the trait is positively selected and rises in frequency. In configuration 2, the preference is initially too rare for the trait to be positively selected, and so the trait allele decreases in frequency. **A.** Without mutation at the trait locus, the trait rapidly rises to fixation in configuration 1, and decreases to extinction in configuration 2. As a correlated response, the preference allele rises in frequency in configuration 1 and decreases in frequency in configuration 2, but the elimination of trait variation among males causes the preference to then stagnate in both cases. **B.** With mutation at the trait locus, variation for the trait is never eliminated, and persistent frequency change of the preference—including to fixation or extinction—becomes possible. Parameters: *α* = 3, *s*_*T*_ = 0.2, *h*_*T*_ = *h*_*p*_ = 1/2, *r* = 0.1, *μ* = 0 (**A**) or *μ* = 0.01 (**B**). Starting frequencies: *T*: 5%, *P*: 25% (configuration 1) or 15% (configuration 2), in Hardy-Weinberg and linkage equilibrium.

### Mutation at the trait locus permits long-term spread of the preference

This third feature, the transience of the increase in frequency of the preference, limits analysis of the long-run dynamics of Fisherian sexual selection in two-locus models. Fortunately, there is a simple (and realistic) modification of the model that leads to longer term increases of the preference: mutation at the trait locus. Fig. 2B shows allele frequency trajectories for the same parameter configurations as in Fig. 2A when the two alleles at the trait locus, *t* and *T*, are allowed to mutate to one another. This causes persistent polymorphism at the trait locus, and therefore makes long-term spread of *P* possible (Faria, Varela & Gardner 2018). We include mutation at the trait locus for all further results, and discuss its role in more detail in the Discussion section.

### Two channels of Fisherian selection in diploids

Spread of *P* is an indirect consequence of its being positively associated with *T*—that is, individuals carrying *T* are disproportionately likely also to carry *P*. If mate choice occurs among diploids, two positive associations between *T* and *P* can be used as information by *P*: a male carrying *T* could be disproportionately likely to carry *P* on the same haploid set of chromosomes (i.e., inherited from the same parent) or on the homologous set (Fig. 1). That is, *T* and *P* can be in positive cis-linkage disequilibrium (c-LD; gametic phase disequilibrium) and/or positive trans-linkage disequilibrium (t-LD). These two sources of information represent two channels of Fisherian sexual selection in diploids.

The ‘trans channel’ is based on positive t-LD between the *P* and *T* alleles, which arises each generation because of the action of the mate preference in the previous generation—a disproportionate number of offspring are produced from matings between preference-bearing females and trait-bearing males. The ‘cis channel’ operates because of positive c-LD between *P* and *T*, which builds up by several mechanisms that we discuss later. Importantly, the generation of t-LD is a simple consequence of the action of the mate preference each generation, and thus does not depend on prior c-LD; in contrast, the generation of c-LD does, in part, depend on prior t-LD between the alleles (below).

### Artificial removal of the cis channel demonstrates independent operation of the trans channel

This asymmetry of the causal relationship between c-LD and t-LD allows us to artificially eliminate the cis channel in simulations while keeping the trans channel intact. We achieve this by eliminating c-LD each generation, by manipulating the relative frequencies of the two double heterozygotes, *Pt*/*pT* and *PT*/*pt*—this procedure alters haplotype frequencies but not diploid genotype frequencies. Note that it does not matter whether we remove c-LD before or after viability selection has acted, because, without epistatic selection, c-LD does not build up within generations. (Note too that, because the c-LD underlying the cis channel relies in part on prior t-LD, we cannot permanently eliminate the trans channel without affecting the cis channel.)

Fig. 3A shows example trajectories of the frequencies of *P* and *T* when this procedure is carried out, together with the trajectories that would result were the cis channel left intact. Several points can be observed: First, *P* can increase in frequency even in the absence of c-LD. This demonstrates independent operation of Fisherian sexual selection via the trans channel. Second, when *P* is positively selected, it spreads more slowly than it would were the cis channel intact. As a check, Fig. 3B shows analogous frequency trajectories of *P* and *T* in the haploid model, when c-LD is artificially eliminated each generation. As expected, *P* does not increase in frequency in this case, since selection on *P* in the haploid model operates only through the cis channel.

**Figure 3.**
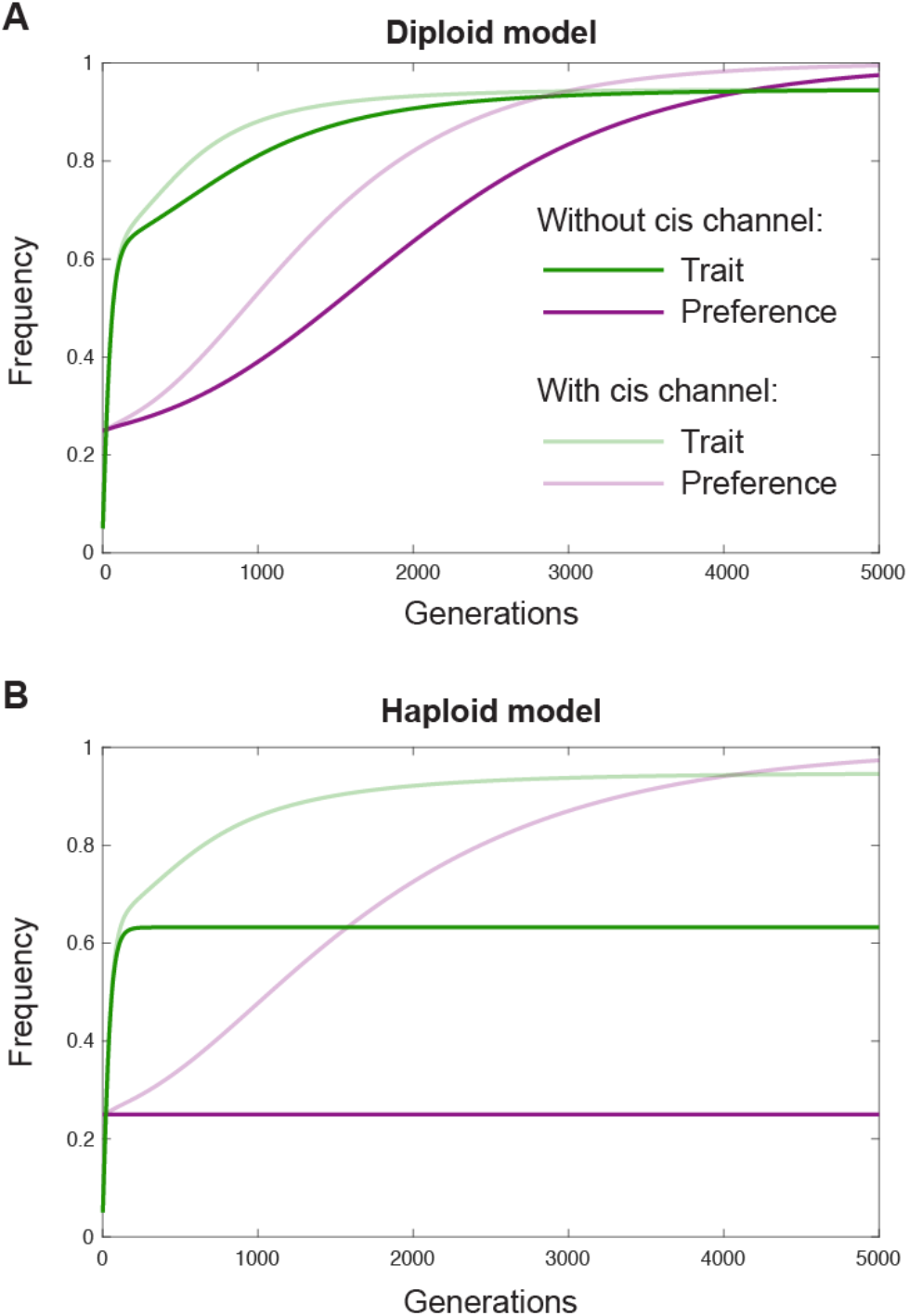
The preference spreads in the diploid model despite elimination of the cis channel. In the text, we describe a procedure to experimentally eliminate cis associations between the preference and trait alleles without changing genotype frequencies. Here, the faint lines indicate allele trajectories when this procedure has not been carried out. **A.** Eliminating the cis channel in the diploid model, we find that the preference can nonetheless spread, indicating that another channel—the trans channel—is operating. Spread of the preference is, however, slower in the absence of the cis channel. **B.** The haploid model, in contrast, depends on cis associations for the spread of the preference, and so the preference cannot spread when we experimentally eliminate these cis associations. Parameters in **A** are the same as in Fig. 2B, configuration 1. Parameters in **B** are the haploid analog of those in 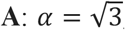, *s*_*T*_ = 0.1, *r* = 0.1, *μ* = 0.01, starting frequencies: *T*: 5%, *P*: 25% in linkage equilibrium.

### The trans channel allows the preference allele to spread with the trait allele even when they are in negative cis-linkage disequilibrium

The presence of an always-positive trans channel in the diploid model implies, in principle, that *P* can spread together with *T* even when they are in negative c-LD, provided the positive trans association is stronger than the negative cis association. While the configurations of the model that we have studied so far lead to positive c-LD, there are some configurations of the model, particularly when the trait allele is fully dominant, that can lead to negative c-LD (discussed below). An example of such a configuration is given in Fig. 4A, where *P* and *T* both increase in frequency despite negative c-LD between them. The spread of the preference here must be due to the trans channel.

**Figure 4.**
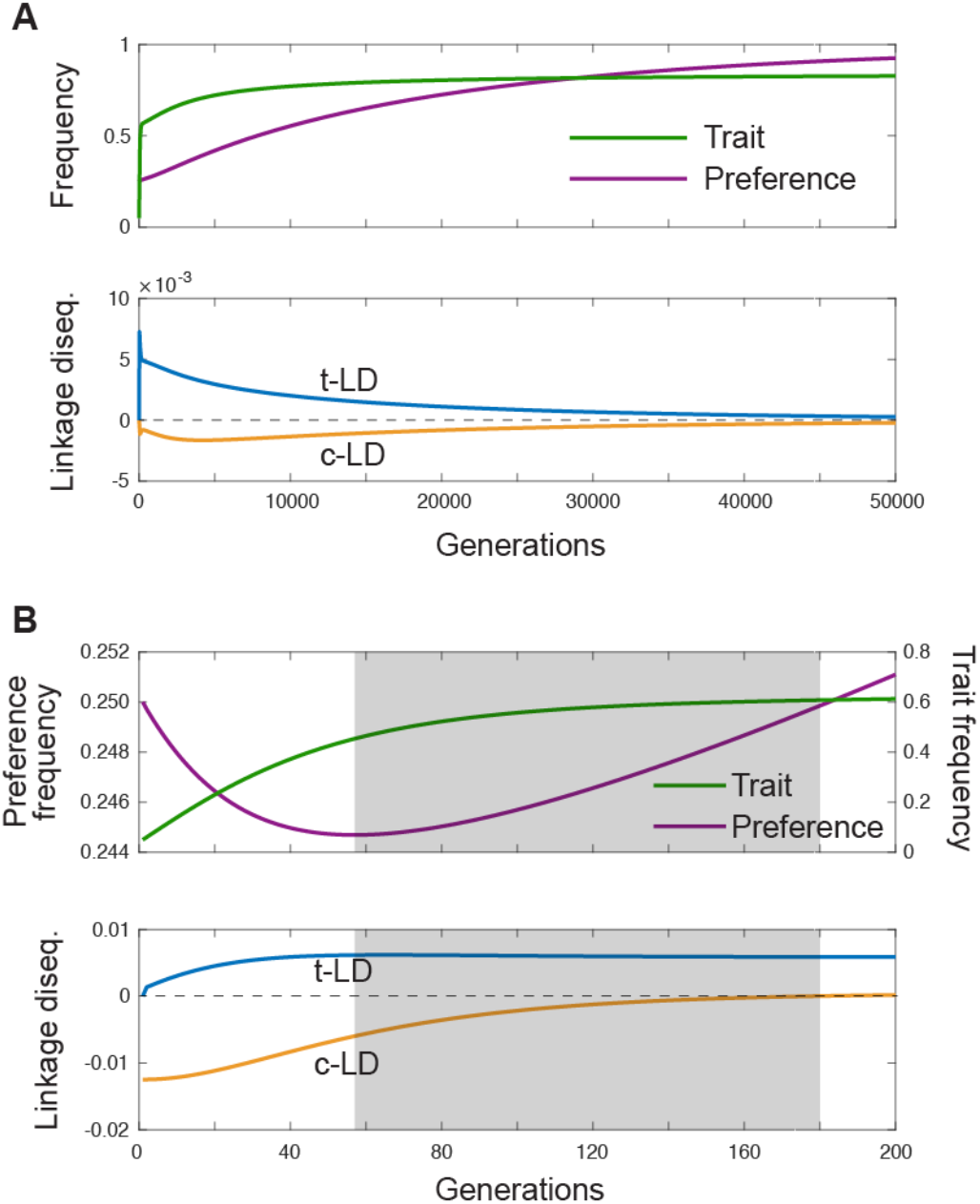
The preference allele can spread in the diploid model despite negative cis-linkage disequilibrium with the trait allele. In the haploid model, co-spread of the trait and preference requires that they be in positive c-LD. In the diploid model, the operation of the trans channel implies that the preference can spread despite being in negative c-LD with the trait. **A.** Negative c-LD is generated naturally in the diploid model when the trait allele is fully dominant and positively selected. Here, the preference allele spreads together with the trait allele despite this negative c-LD, because the positive trans association between the alleles (t-LD) outweighs their negative cis association (c-LD). **B.** c-LD can be forced initially to be negative in our focal case (where the trait allele is co-dominant) by starting the population without the *PT* and *pt* haplotypes, and eliminating recombination. Although positive c-LD is eventually restored in this case, as the *PT* and *pt* haplotypes are generated by mutation, there is a period (shaded grey) where c-LD is negative and the preference and trait are nonetheless co-spreading. Parameters: As in Fig. 2B, configuration 1, but *h*_*T*_ = 1 (**A**), or *r* = 0 and the starting state excludes the *pt* or *PT* haplotypes (**B**). In **B**, the starting allele frequencies are still *T*: 5%, *P*: 25%.

An experimental demonstration that *P* can spread despite being in negative c-LD with *T* can also be achieved in our baseline case of co-dominance of both alleles (*h*_*T*_ = *h*_*p*_ = 1/2), by beginning simulations without the *PT* haplotype. We further assume that there is no recombination between the two loci, so that the *PT* haplotype can only be restored slowly by mutation (note that restoration of the *PT* haplotype is required because, were it permanently absent, *P* could never increase in frequency). c-LD is thus forced initially to be negative (with *q*_*PT*_ ≈ 0, c-LD = *q*_*PT*_ *q*_*pt*_ − *q*_*Pt*_*q*_*pT*_ ≈ −*q*_*Pt*_*q*_*pT*_ < 0, where *q*(·) is the frequency of the subscripted haplotype among diploids). Fig. 4B shows an example in this case where *P* spreads during a period when c-LD is still negative. The spread of the preference while c-LD is negative must be due to the trans channel.

## Relative importance of the trans and cis channels

### Trans associations of trait and preference are more informative when linkage is tight

The results above demonstrate that sexual selection on female mate preferences in diploids operates through two channels, the trans channel and the cis channel. Under the greenbeard interpretation of Fisherian sexual selection, these constitute, for a copy of *P* acting in a female, two channels of information about the likelihood that a trait-displaying male carries *P* too. Therefore, a natural way to compare the relative importance of the two channels is to compare how informative they are in this regard. That is, considering a given copy of *T* in a male, what are the (conditional) probabilities that he carries *P* in trans, and in cis, with respect to the focal trait allele? With no association, this probability would simply be the frequency of *P*, and so we are interested in how much the two conditional probabilities exceed this baseline value. The ratio of the excess of these two probabilities over the baseline value gives the relative importance of the trans channel over the cis channel, and is in fact equal to the ratio of t-LD and c-LD.

Fig. 5A shows the trajectory of this ratio in the diploid model, for various recombination rates. There are several key observations: First, the trans channel is always at least as important as the cis channel (among the configurations we have tested). Second, the relative importance of the trans channel over the cis channel increases as the recombination rate between the preference and trait loci decreases. When recombination is free, the cis and trans channels are of approximately equal importance, consistent with analytical results that c-LD and t-LD are equal in the quasi-linkage equilibrium (QLE) limit of the diploid model, where recombination dominates selection (Kirkpatrick, Johnson & Barton 2002 [Eq. 56]; Greenspoon & Otto 2009 [Eq. 6]). When linkage is tight, the trans channel becomes much more important than the cis channel—e.g., as much as 300 times more important for the parameters used in Fig. 5.

**Figure 5.**
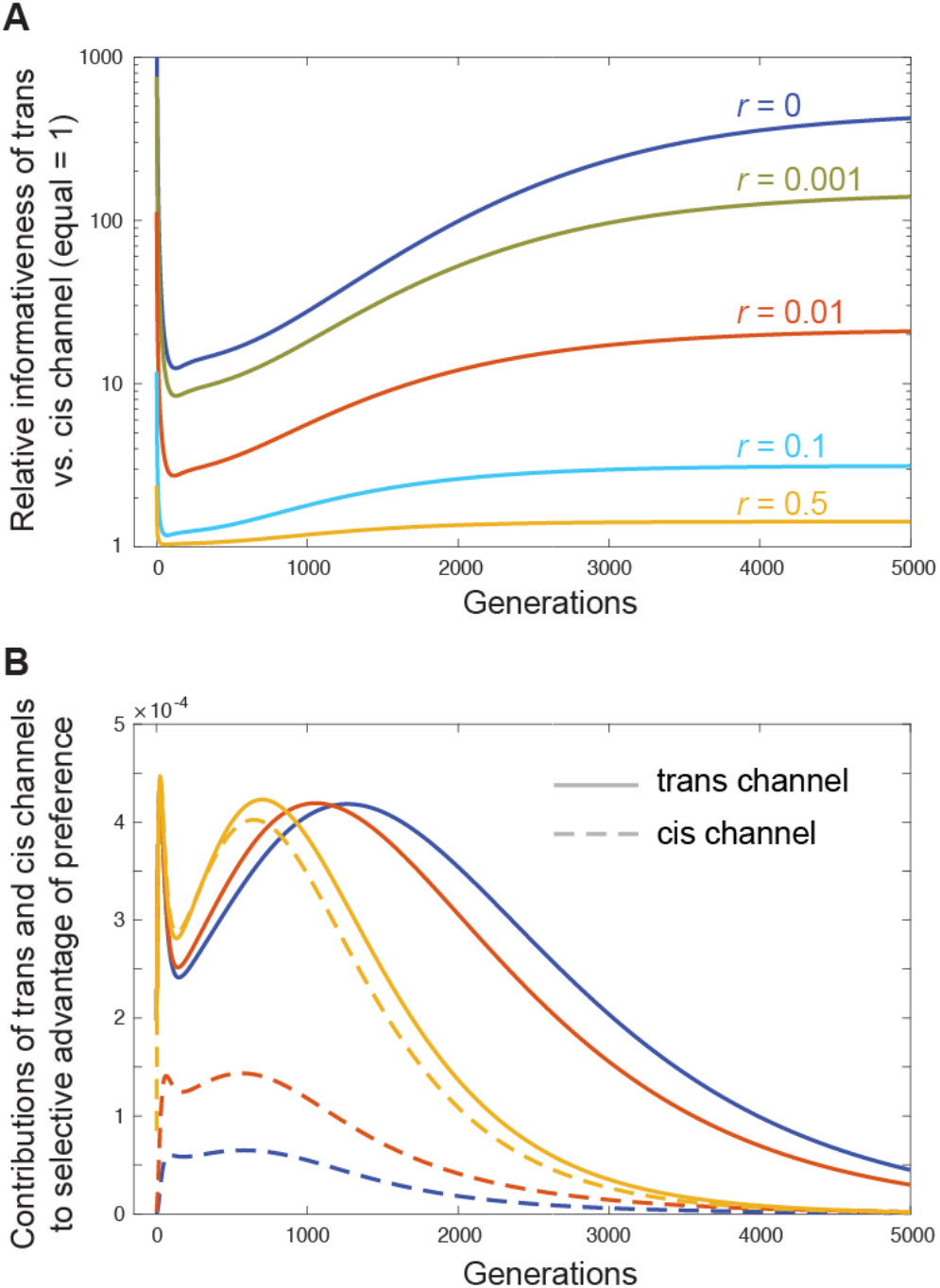
Relative importance of the trans channel versus the cis channel. **A.** The ratio of t-LD to c-LD, for various recombination rates. Under the greenbeard interpretation, this is the ratio of the two channels’ informativeness to a preference allele in a female that a trait-displaying male is likely to carry the preference allele. **B.** By temporarily disabling the trans channel each generation (procedure described in text), we can observe how much of the frequency increase of the preference allele is due to the cis channel, and therefore also how much is due to the trans channel. These contributions are plotted for selected recombination rates from **A**. Both **A** and **B** reveal that the relative importance of the trans channel increases as the recombination fraction between the two loci decreases. Parameters: As in Fig. 2B, configuration 1, except for variable *r*.

### The trans channel contributes more to the spread of the preference when linkage is tight

A second way to assay the relative importance of the trans and cis channels is to ask how much of the selective advantage of the preference is due to each—that is, to decompose frequency increases of *P* into a component due to the trans channel and a component due to the cis channel. We achieve this in our simulations in the following way. Each generation *g*, we construct two populations of zygotes: a true population taking into account the action of the mate preference in generation *g*– 1, and a hypothetical population assuming random mating in *g*– 1. We then calculate the frequency change of *P* from zygotes in *g* to zygotes in *g* + 1 for both populations, taking into account the action of the mate preference among adults in *g*. Because the hypothetical population in *g* has no trans association between *P* and *T* (its parents mated randomly), any increase of *P* between *g* and *g* + 1 must be due solely to the cis channel. Therefore, the difference between the frequency increases of *P* in the true and hypothetical populations identifies the contribution of the trans channel. (Note that this procedure does not permanently remove the trans channel—as pointed out earlier, this would permanently affect the cis channel as well. Instead, this procedure temporarily removes the trans channel each generation, but restores it for the production of true genotype frequencies in the next generation.)

Fig. 5B shows some examples trajectories of the contributions of the trans and cis channels to frequency increases of *P*. Consistent with the conditional probability results (Fig. 5A), we find that the trans channel always contributes at least as much to the selective advantage of the preference as the cis channel does, and that the relative contribution of the trans channel increases as the recombination rate between the preference and trait loci decreases. Thus, when recombination is free, the contributions of the trans and cis channels are approximately equal, consistent with previous analytical results in the QLE limit (Kirkpatrick, Johnson & Barton 2002; Greenspoon & Otto 2009), but when recombination is rare, the trans channel’s contribution dominates.

## Multiple mechanisms for the generation of cis-linkage disequilibrium

### The recombination mechanism

The simplest mechanism by which c-LD is generated in Fisherian sexual selection is by recombination. The mate preference acting in one generation causes the *P* and *T* alleles to be disproportionately associated in trans in the next generation, thus giving them disproportionate opportunities to recombine onto the same background in gametogenesis. When recombination occurs at a higher rate, this ‘recombination mechanism’ is stronger, and thus positive c-LD can build up faster. The recombination mechanism promotes positive c-LD quite generally in models of Fisherian sexual selection, e.g., across different dominance values of the trait and preference alleles.

However, a surprising finding in the above analyses is that, in the diploid model (but not the haploid model), positive c-LD is generated in the absence of recombination when the trait is co-dominant, starting from a state of linkage equilibrium (Fig. 7). This is why the trans channel is only finitely more important than the cis channel when recombination is absent in the diploid model (Fig. 5), and points to mechanisms other than recombination for the generation of c-LD.

### The dominance mechanism

A second mechanism for the generation of c-LD is exclusive to the diploid model, and can operate even in the absence of recombination. It arises from differences in the transmission rates of the various haplotypes, caused by an interaction between (i) their non-random association with other haplotypes in diploid males, and (ii) the dominance of the trait allele *T*.

The operation of this ‘dominance mechanism’ is easiest to see in the case where recombination is absent, the system begins in linkage equilibrium, and the preference allele *P* starts at a sufficiently high frequency that trait-bearing males are positively selected. In this case, there is no LD in the gametes of first-generation males or females, nor is there c-LD in second-generation zygotes. However, a consequence of the mate preference in the first generation is that madumnal (maternally-derived) *PT* and *Pt* haplotypes in second-generation males are disproportionately associated with padumnal (paternally-derived) *T* alleles, while madumnal *pT* and *pt* haplotypes are not.

If *T* is fully recessive, these associations give *PT* haplotypes in second-generation males (which disproportionately reside in *TT* males) a transmission advantage over *pT* haplotypes, but give *Pt* haplotypes (which disproportionately reside in *Tt* males) no advantage over *pt* haplotypes. Thus, while haplotype frequencies in second-generation males obey *q*_*PT*_/*q*_*pT*_ = *q*_*Pt*_/*q*_*pt*_ (no c-LD), haplotype frequencies in the successful sperm of these males obey *q*_*PT*_/*q*_*pT*_ > *q*_*Pt*_/*q*_*pt*_ (positive c-LD). On the other hand, if *T* is fully dominant, then the haplotype associations in males give *PT* haplotypes no advantage over *pT* haplotypes (since there is no advantage to being associated with another *T* allele), but give *Pt* haplotypes an advantage over *pt* haplotypes, so that *q*_*PT*_/*q*_*pT*_ < *q*_*Pt*_/*q*_*pt*_ among sperm (negative c-LD). For some intermediate level of dominance, *q*_*PT*_/*q*_*pT*_ = *q*_*Pt*_/*q*_*pt*_ among sperm (no c-LD generated by the dominance mechanism).

Thus, the ‘dominance channel’ generates positive c-LD when *T* is recessive, negative c-LD when *T* is dominant, and no c-LD when *T* has a certain intermediate dominance (Fig. 7). Though we have focused, for clarity, on the first few generations in the case where recombination is absent, the logic above relies only on the fact that madumnal *PT* and *Pt* haplotypes associate disproportionately with padumnal *T* alleles—the dominance of *T* then governs the transmission advantages these associations confer on the *PT* and *Pt* haplotypes relative to the *pT* and *pt* haplotypes, respectively, and thus the effect on c-LD.

It is clear that the dominance mechanism cannot operate in the haploid model, because *PT* and *pT* males are equally fit, and so these haplotypes have equal transmission rates in sperm; the same holds for the *Pt* and *pt* haplotypes.

### The sexual admixture mechanism

A third mechanism for the generation of c-LD derives from sex-specific selection at both loci. If *P* and *T* are associated (either in trans or in cis), then male-specific viability selection on *T* implies indirect male-specific selection on *P*, causing the frequencies of both alleles to differ between sperm and eggs. Allele frequency differences at multiple loci in sperm and eggs create c-LD upon sexual admixture (Úbeda, Haig & Patten 2010).

This argument applies equally to the haploid and diploid models, so why is c-LD generated in the absence of recombination in the diploid model but not the haploid model? The ‘sexual admixture mechanism’ for generating c-LD requires a prior association between *P* and *T*, so that male-specific selection on the trait changes the frequencies of both alleles in sperm relative to eggs. In the haploid model, the only possible prior association is c-LD, and so the sexual admixture mechanism cannot generate c-LD without initial c-LD (Fig. 6). In the diploid model, even in the absence of recombination, *P* and *T* immediately come into a trans association by the action of the mate preference. Thus, selection on the trait in males causes the frequencies of both alleles to differ between sperm and eggs, leading to c-LD by the sexual admixture mechanism.

**Figure 6.**
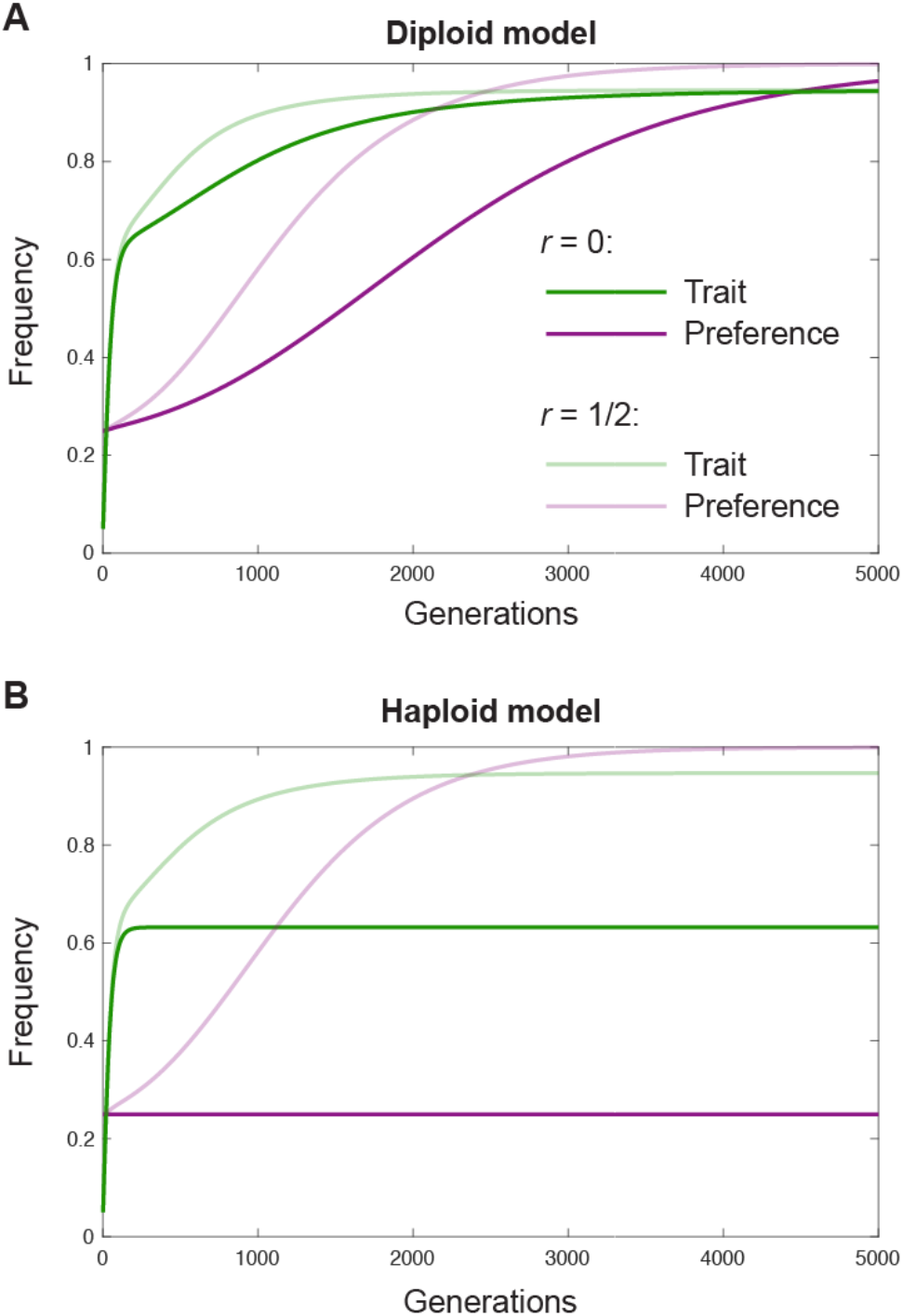
The preference can spread in the absence of recombination in the diploid model, but not the haploid model. The population begins with cis-linkage equilibrium between the two loci. This linkage equilibrium is maintained in the haploid model (**B**), and, because the preference can only increase by the cis channel in the haploid model, this maintenance of c-LD implies that the preference stagnates. In the diploid model (**B**), the preference increases in the absence of recombination, at first because of the operation of the trans channel, but subsequently also because of the operation of the cis channel as c-LD builds up for reasons specific to the diploid model that are explored in the text. Parameters: As in Fig. 2B, configuration 1 (**A**), and Fig. 3B (**B**), except for *r*.

**Figure 7.**
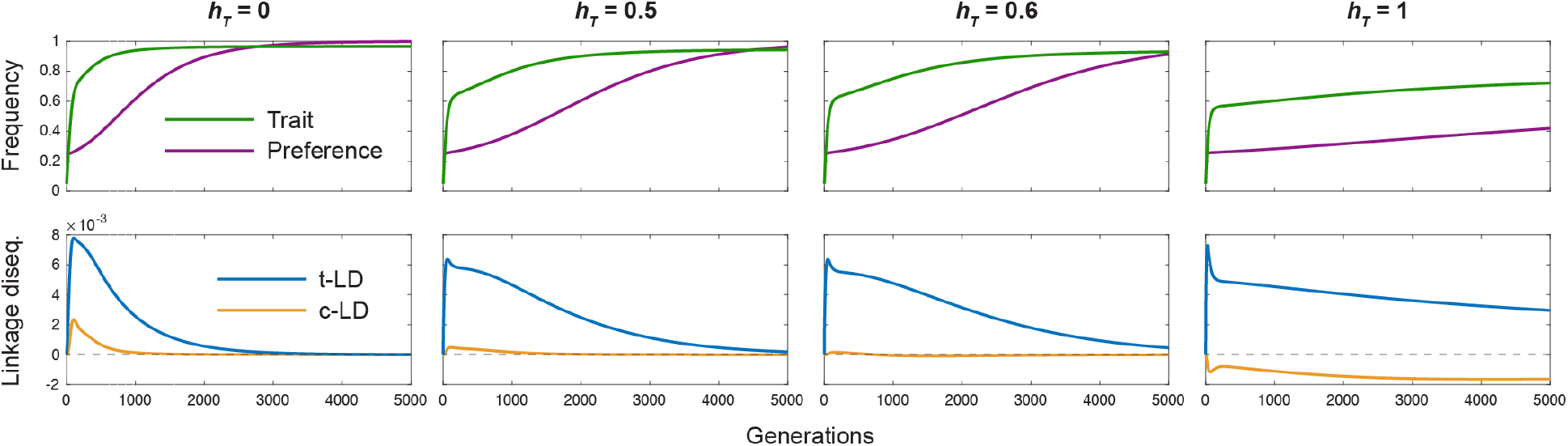
The sign of cis-linkage disequilibrium can depend on dominance of the trait allele. Here, recombination is assumed to be absent, so that the only sources of c-LD in the diploid model are the ‘dominance mechanism’ and, reinforcing this, the ‘sexual admixture’ mechanism. c-LD is positive when the trait is recessive, weakly positive when the trait is co-dominant, approximately zero for an intermediate value of dominance of the trait (*h*_*T*_ = 0.6), and negative when the trait is fully dominant (although note that the preference nonetheless spreads in this case, because of the operation of the trans channel). Parameters: As in Fig. 2B, configuration 1, except for *h*_*T*_.

In the haploid model, if c-LD is initially generated for other reasons (e.g., by the recombination mechanism, or by random drift), then the ‘sexual admixture’ mechanism does operate. Eq. (1c) in Kirkpatrick (1982) provides a general expression for the total change in c-LD from one generation to the next in the haploid model. In an Appendix, we isolate from this expression the component due to the sexual admixture mechanism, in the case where *r* = 0.

Notice that the effect of the sexual admixture mechanism on c-LD depends on the sign of the pre-existing association between *P* and *T*. If there is a net positive association between the alleles (e.g., if t-LD and c-LD are both positive, or if negative c-LD arising from the dominance mechanism is outweighed by positive t-LD arising directly by the action of the mate preference), then they will change in frequency in the same direction in sperm, and so the sexual admixture mechanism makes a positive contribution to c-LD in the next generation (Úbeda, Haig & Patten 2010). If there is a net negative association between the alleles (e.g., if negative c-LD arising from the dominance mechanism outweighs the positive t-LD generated by non-random mating), then they will change in frequency in opposite directions in sperm, and so the sexual admixture mechanism makes a negative contribution to c-LD. Thus, the sexual admixture mechanism only reinforces pre-existing associations between the alleles.

## Discussion

The life cycle of all sexual organisms involves alternation between a haploid and a diploid phase (Mable & Otto 1998). Sexual selection can operate in either phase, leading to separate haploid and diploid models of Fisherian sexual selection. Classical work characterized the population dynamics of the two-locus haploid model (Kirkpatrick 1982); later work identified differences in the dynamics of diploid model, particularly relating to the structure of stable equilibrium sets (e.g., Gomulkiewicz & Hastings 1990; Heisler & Curtsinger 1990). However, as noted by Barton & Turelli (1991), many of these differences are due not to the action of sexual selection in the diploid phase *per se*, but rather to the additional parametric complexity of the diploid model. Here, we have identified a fundamental conceptual difference between the haploid and diploid models of Fisherian sexual selection.

In doing so, we have made use of the greenbeard interpretation of Fisherian sexual selection (Dawkins 1986; Pizzari & Gardner 2012; Faria, Varela & Gardner 2018). Under this interpretation, the trait displayed by a male is used by a preference allele in a female as information that the male probably carries the preference allele. The preference allele in the female induces her to mate with the male, thus likely conferring a mating advantage on a copy of itself in the male. This interpretation naturally raises the question: What is the mechanism by which a male’s display of the trait is informative that he likely carries the preference allele? That is, what is the nature of the association between the trait and preference alleles?

If sexual selection operates in the haploid phase, there is only one way in which the trait and preference alleles can be associated in males: they must be in cis-linkage disequilibrium (c-LD; gametic phase disequilibrium). However, if sexual selection occurs in the diploid phase, there are two ways in which the trait and preference alleles can be associated: they can be in c-LD or in trans-linkage disequilibrium (t-LD). These represent two distinct ways in which the trait displayed by a diploid male is informative that he carries the preference allele, and they thus represent two distinct channels of Fisherian sexual selection in diploids. We have named these two channels the ‘trans channel’ and the ‘cis channel’.

We have formally demonstrated the independent action of the trans channel in the diploid model by comparing, in simulations, the population dynamics of the trait and preference alleles when the cis channel is kept open and when it is switched off (by the artificial removal of c-LD each generation). When the cis channel is switched off in the haploid model, the preference is unable to increase in frequency (Fig. 3B). In contrast, in the diploid channel, the preference can still increase in frequency when the cis channel is switched off (Fig. 3A). This increase in frequency in the diploid model is due to the independent operation of the trans channel.

A subtlety is that, in both the haploid and diploid models of Fisherian sexual selection, the generation of c-LD (and thus the operation of the cis channel) depends on repeated trans associations of the preference and trait alleles in the diploid phase of the lifecycle. The key difference between the haploid and diploid models, in terms of operation of the trans channel, is that sexual selection acts in the diploid phase in the ‘diploid model’ but not in the ‘haploid model’. Therefore, under the greenbeard interpretation, the trans association is itself informative to the preference in the diploid model, in addition to its fundamental role in generating c-LD; this is not the case in the haploid model.

Recombination has played an important role in our findings. When the recombination rate between preference and trait loci is low, c-LD builds up more slowly, and to a lower level, than when the recombination rate is high. As a result, the trans channel—which is insensitive to the recombination rate, being a direct consequence of the action of the mate preference in the previous generation—is more important than the cis channel when the recombination rate is low (Fig. 5). An implication of this result is that lowering the recombination rate between the trait and preference loci severely slows down Fisherian sexual selection in the haploid model, where cis-linkage disequilibrium is the only mechanism by which the preference can increase in frequency, but has a less severe effect in the diploid model, with its ever-present trans channel (Fig. 8).

**Figure 8.**
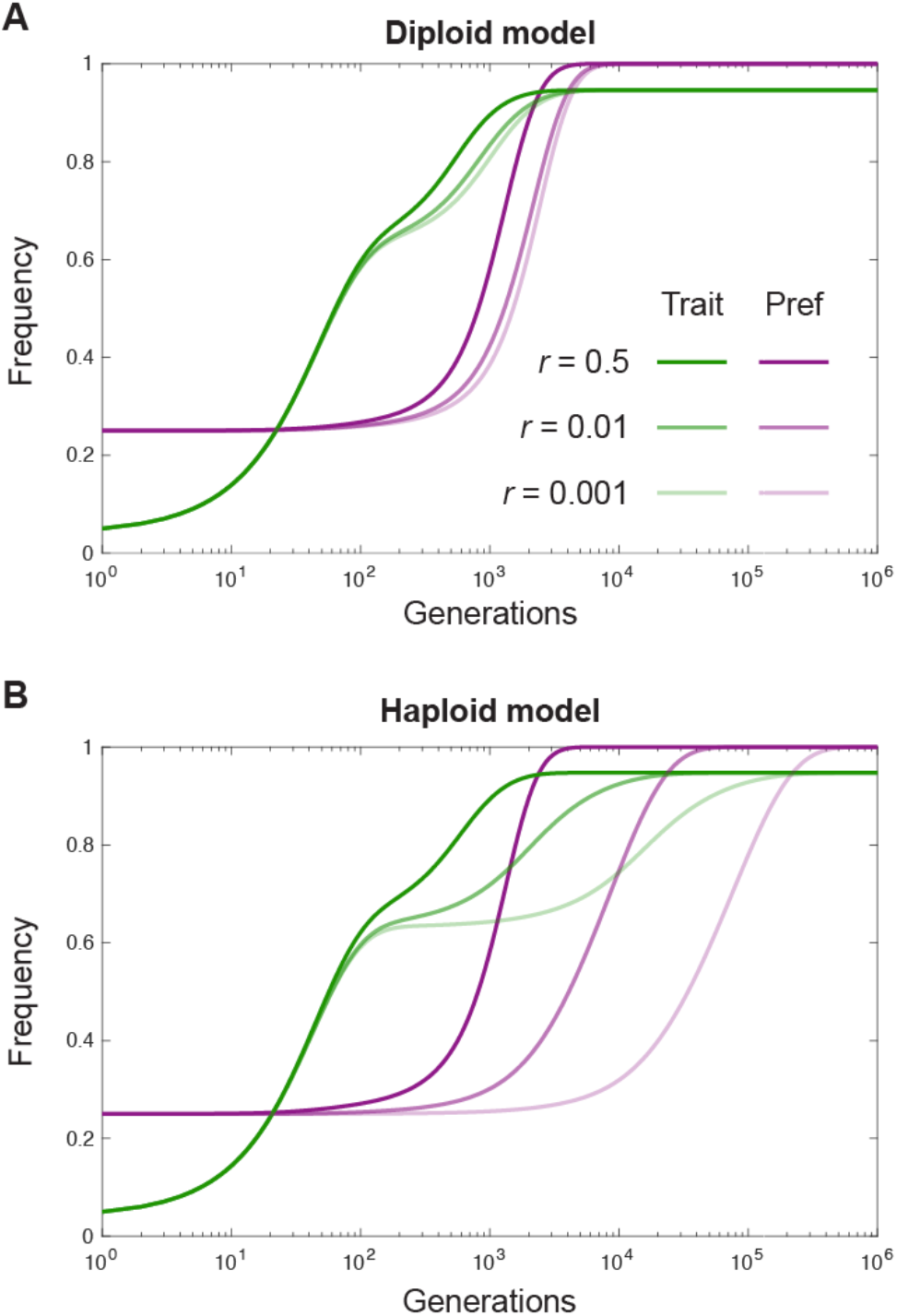
Tight linkage severely slows down Fisherian sexual selection in the haploid model, but not the diploid model. Trajectories of the trait and preference alleles in the diploid (**A**) and haploid (**B**) models, for various recombination rates between the trait and preference loci. The same equilibrium is ultimately reached in each case, but the number of generations required to approach the equilibrium is very large for low recombination rates in the haploid model (note the logarithmic scale of the time axis). This effect is less severe in the diploid model, owing to the ever-present, recombination-insensitive trans channel. Parameters: As in Fig. 2B, configuration 1 (**A**), and Fig. 3B (**B**), except for *r*.

Nonetheless, the above considerations, together with the combinatorial fact that, in most species, the vast majority of locus pairs lie on separate chromosomes (Crow 1988, Veller, Kleckner & Nowak 2019), suggest that two-locus Fisherian systems should usually involve unlinked loci. There are relatively few instances of mate choice for which the genetic bases of both preference and trait have been localized (Chenoweth & Blows 2006). However, in some of these, loci contributing to variation in preference and trait have been found to be tightly linked (e.g., Kronforst et al. 2007, Shaw & Lesnick 2009, Bay et al. 2017, Xu & Shaw 2019). In these cases, our results suggest that the operation of the trans channel was vital for the establishment of these preference-trait gene pairs (the examples all involve sexual selection in diploids), unless the preference and trait alleles are sufficiently tightly linked and appeared already at high cis-linkage disequilibrium and at a sufficient frequency to initiate the Fisherian process.

Interestingly, the examples of tightly linked preference and trait genes cited above are all in clades with recent species radiations and ongoing hybridization. In such cases, tight linkage of preference and trait genes (or pleiotropy of a single gene) might be beneficial in preventing their dissociation in the face of introgression (Xu & Shaw 2019). If so, this would be an instance of a more general evolutionary phenomenon where introgression favors tight linkage of co-adapted gene complexes (Kirkpatrick & Barton 2006). From a theoretical perspective, there are two scenarios: (i) Sexual selection on the preference gene is weaker when it is tightly linked to the trait gene, so that such complexes fix more slowly and less frequently (consistent with our results); however, when they do, they are more robust in the face of introgression and therefore persist while complexes with looser linkage do not; (ii) Ongoing introgression in fact causes sexual selection on a preference gene to be stronger when it is tightly linked to the trait gene, so that tightly linked complexes fix more rapidly and more frequently. More modelling of Fisherian sexual selection with introgression (e.g., Servedio & Bürger 2018) is required to understand the relative likelihood of these two possibilities. Note that the results of such models are likely to be sensitive to whether sexual selection occurs in the haploid or the diploid phase, since, as we have found, the trans channel dominates in diploid sexual selection when linkage is tight, but does not exist in haploid sexual selection.

We have started all of our simulations with the preference and trait alleles in Hardy-Weinberg and cis-linkage equilibrium. From such an initial state, when recombination between the loci is absent, only weak c-LD between the alleles can build up (and in the haploid model, none at all). Thus, the trans channel, driven directly by assortative mating, dominates. However, there are instances where we expect Fisherian sexual selection within a population to begin from a state where the preference and trait alleles are already in strong c-LD. For example, if the preference and trait loci lie within a chromosomal segment that is inverted in one population relative to the other, then introgression of preference and trait alleles from a population where they are fixed into the other where they are absent would constitute an injection of preference and trait alleles in perfect c-LD. Because they reside in an inversion in the recipient population, recombination cannot dissociate them, and their c-LD is maintained. Notice, however, that the preference-trait haplotype must be introgressed at a sufficiently high frequency that the mating advantage conferred on the introgressed trait outweighs its viability disadvantage. If so, the cis-channel would dominate in the resulting positive Fisherian sexual selection of the introgressed preference. This is in sharp contrast to our results concerning the importance of the cis and trans channels when recombination is absent and the preference and trait alleles start in linkage equilibrium, and thus highlights the role of initial linkage in how the Fisherian process plays out.

Although we have focused on autosomally encoded preferences and traits in diploids, our conceptual distinction between the cis and trans channels generalizes easily to other systems. Consider X-linked preference and trait alleles in a male-heterogametic (XX/XY) species. In this case, although females are diploid at the preference and trait loci, males are effectively haploid—under the greenbeard interpretation, only the cis channel provides information to a preference allele in a female that a trait-bearing male is likely also to carry the preference allele (because, if he does, both his preference and trait allele must lie on the same X chromosome). In contrast, Z-linked traits in female-heterogametic (ZW/ZZ) species are informative to Z-linked preferences via both the cis and trans channels, because males are diploid for the Z. This suggests that tightly-linked preference and trait alleles should be much more common on Z chromosomes than on X chromosomes, because tight linkage will severely slow down their co-spread in XX/XY species (where only the recombination-sensitive cis channel is operating) but not in ZW/ZZ species (where the recombination-insensitive trans channel operates too).

The situation is slightly more complicated when only the trait locus is sex-linked. An X-linked trait allele carried by a male was necessarily inherited from his mother. Thus, even though a trans association between the preference and trait alleles is technically possible in this case (because of diploidy at the preference locus in males), the trans channel, which functions in the autosomal diploid model because trait-bearing males are disproportionately likely to have inherited the trait allele from their fathers and the preference allele from their mothers, in fact does not operate here. In contrast, both the cis and trans channels operate when the trait is Z-linked and the preference is autosomal, because each male inherits a Z from both his mother and his father. However, because recombination is free when the trait is sex-linked but the preference is autosomal, the absence of the trans channel in XX/XY species in this case is not expected to be influential.

The situation for haplodiploid species is the same as the case where both the preference and trait are X-linked, with the additional point that the preference and trait loci can lie on separate chromosomes in haplodiploids—an important point, given that only the cis channel operates in this case, so that free recombination between the preference and trait loci (in females) is very helpful. Finally, we have considered the case of female mating preferences for male traits, but all of our results apply, *mutatis mutandis*, to male mating preferences for female traits, which are now recognized to play an important role in many systems (Edward & Chapman 2011, Fitzpatrick & Servedio 2018). The results for autosomal traits and preferences are the same as those for female choice, while the statements above for XX/XY and ZW/ZZ species reverse.

When the only forces affecting the frequencies of the preference and trait alleles are viability selection and the mate preference, frequency changes of the preference allele are typically transient, as the trait is rapidly driven either to fixation or extinction (Fig. 2A). A simple augmentation of the model that allows for longer-term frequency changes of the preference allele is mutation at the trait locus (Fig. 2B). The effect of mutation can be understood from the case where the preference is initially sufficiently common that the trait allele *T* is positively selected. *T* rapidly spreads to a mutation-selection balance near its fixation boundary. At this mutation-selection balance, the fitness advantage of *T* (owing to the common mate preference) is a force pushing it up in frequency, while mutation is a force pushing it down in frequency (since *T* is much more common than *t*). Because the preference allele *P* is in a positive association with *T*, it benefits indirectly from *T*’s overall fitness advantage without suffering from the net mutational loss of *T* alleles. *P* thus increases in frequency as long as polymorphism is maintained at the trait locus.

We have focused on the ‘Fisherian’ case where the only selection acting on the mate preference is indirect, as a correlated response to direct selection on the trait. However, discriminatory mate choice is often associated with direct fitness costs, reducing the number of offspring that choosy females have (Kirkpatrick 1987, Pomiankowski 1987). The persistent indirect selective advantage to *P* that is generated by mutation at the trait locus allows *P* to spread even when it carries a direct fitness cost, provided that this cost is not too large (Fig. 9) This is impossible in models without trait mutation—either two-locus or quantitative— where direct costs of the preference lead to its eventual extinction (Kirkpatrick 1985; Pomiankowski 1987; Pomiankowski, Iwasa & Nee 1991). Importantly, in the two-locus model, mutation at the trait locus can lead to persistent spread of an intrinsically costly preference no matter the relative mutation rates from *t* → *T* and *T* → *t* (as long as the latter is not zero). In particular, a costly preference can spread even when there is a substantial bias in favor of *t* → *T* mutations (Fig. 9). This is in contrast to quantitative genetic models of Fisher’s process, where a costly preference can spread in the long term only if there is a mutational bias away from the preferred trait (Pomiankowski, Iwasa & Nee 1991).

**Figure 9.**
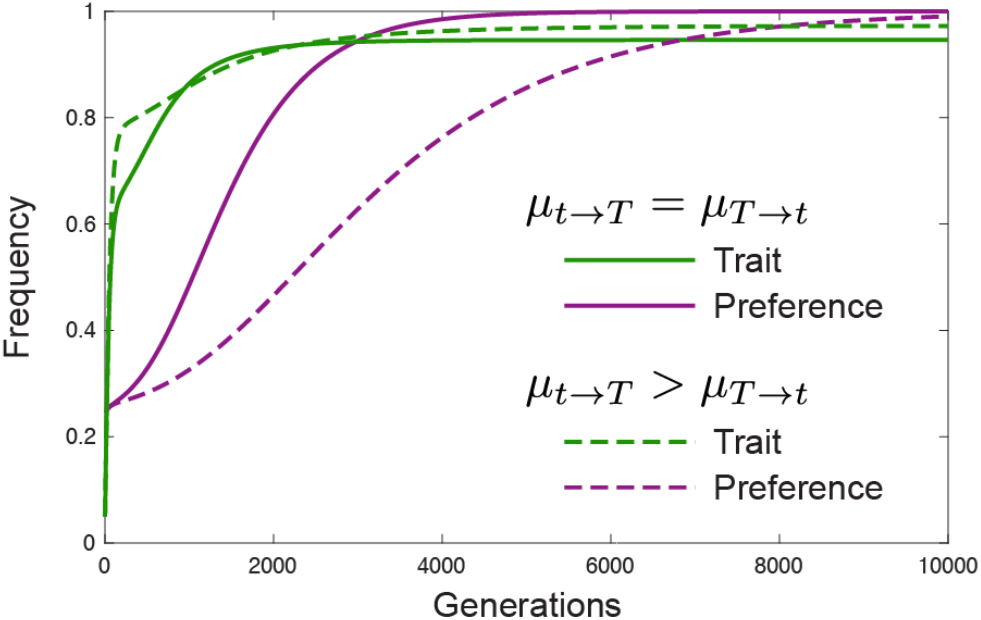
Mutation at the trait locus allows an intrinsically costly preference to spread. A positive *T* → *t* mutation rate generates a persistent selective advantage for the preference, allowing the preference allele *P* to spread even when it is intrinsically costly, with *Pp* and *PP* females having reduced viability or fecundity 1– *h*_*p*_ *s*_*P*_ and 1– *s*_*P*_ respectively, relative to *pp* females. The preference can spread even when the positive *T* → *t* mutation rate is smaller than the *t* → *T* mutation rate, in contrast to quantitative genetic models where mutation away from the preferred trait must be more common than the reverse for a costly preference to spread in the long term. Parameters: As in Fig. 2B, configuration 1, with *s*_*P*_ = 0.001, *h*_*p*_ = 1/2 and with *μ*_*T*→*t*_ = 0.01 and *μ*_*t*→*T*_ = 0.01 (symmetric case) or *μ*_*T*→*t*_ = 0.005 and *μ*_*t*→*T*_ = 0.01 (asymmetric case).

## Acknowledgments

We are grateful to Graham Coop and Mark Kirkpatrick for helpful discussions.

## Appendix: Contribution of the sexual admixture mechanism to LD in the haploid model

We assume that natural and sexual selection act in the haploid phase, and that there is no recombination in the diploid phase. Order the four haploid genotypes *PT*, *Pt*, *pT*, *pt*, and suppose that their frequencies before viability selection are *x*_1_, *x*_2_, *x*_3_, *x*_4_. These frequencies are equal for females and males. The frequency of the *P* allele is *x*_*P*_ = *x*_1_ + *x*_2_ and the frequency of the *T* allele is *x*_*T*_ = *x*_1_ + *x*_3_. The value of linkage disequilibrium is *D* = *x*_1_*x*_4_ − *x*_2_*x*_3_ = *x*_1_ − *x*_*P*_ *x*_*T*_.

Suppose that males with the *T* allele have a relative fitness advantage *S*—this is a function of the strength of viability selection against such males (*s*), the strength of the mating preference (*α*), and the proportion of females exhibiting the preference (*x*_*P*_)—see below.

From Eq. (2.5) of Úbeda, Haig & Patten (2011), corrected for a typo, LD in the next generation is

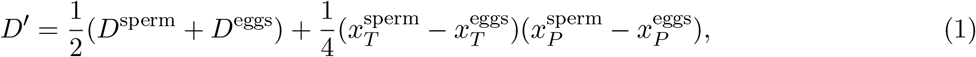

where (·)^sperm^ and (·)^eggs^ are values in successful sperm and eggs.^1^ The first term in Eq. (1) is the contribution to next generation’s LD from LD in the previous generation, while the second term is the contribution from allele frequencies at both loci being different in sperm and egg (the ‘sexual admixture mechanism’).

Because there is no selection among female genotypes, *D*^eggs^ = *D*, 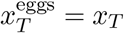, and 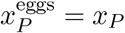. Therefore, Eq. (1) becomes

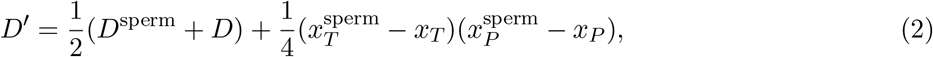

which can be rearranged to become

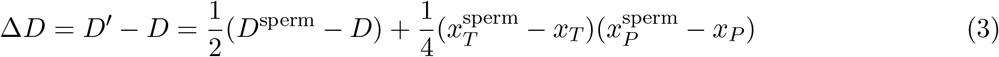

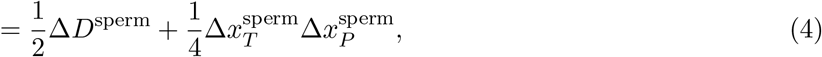

where Δ(·)^sperm^ signifies a difference in values between males and their successful sperm. Among successful sperm:

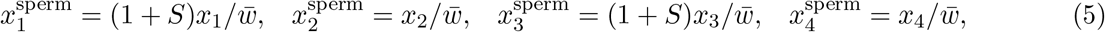

where 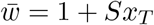. From these,

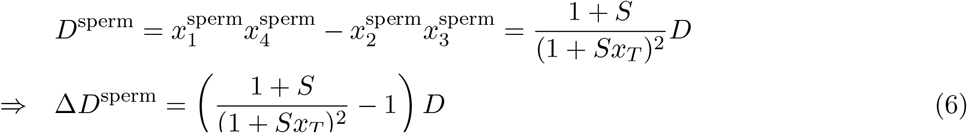

and

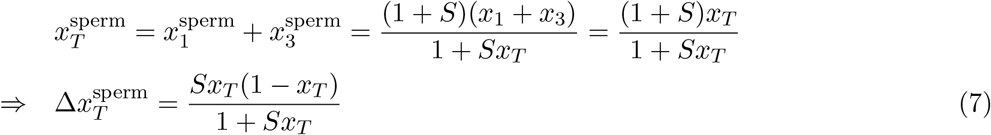

and

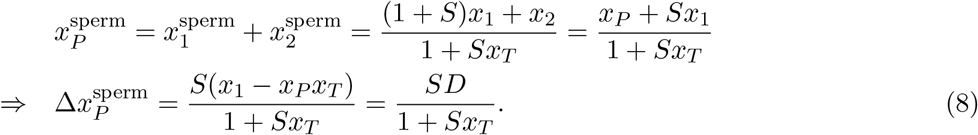

Substituting into Eq. (4),

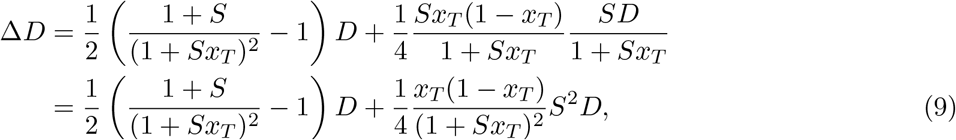

where the second term is the contribution of the sexual admixture mechanism.

To show that this is equivalent to Eq. 1c in Kirkpatrick (1982) in the case where *r* = 0, and thus to identify in Kirkpatrick’s expression the contribution of the sexual admixture mechanism, we solve for *S* as a function of the basic components of selection in the model, with the notational substitutions *α* for Kirkpatrick’s *a*_2_, *x*_*T*_ for his *t*_2_ (so that his *t*_1_ = 1 − *x*_*T*_), and *x*_*P*_ for his *p*_2_ (so that his *p*_1_ = 1 − *x*_*P*_). Then, using the frequencies of the haplotypes *Pt* and *pT* among males before selection, and among successful sperm after selection (Kirkpatrick’s Table 1), we find

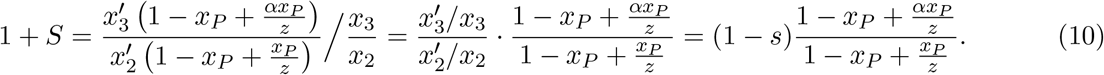

Thus, in the notation of Kirkpatrick’s Eq. 1, *S* = *B*/*A* − 1. Moreover, from our Eq. (7), 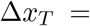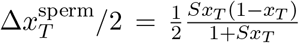, which we equate with Kirkpatrick’s Eq. 1c to find 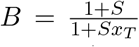, which gives 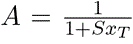. From these, we find the identities 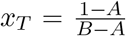, 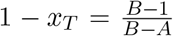, and 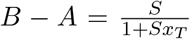. From these, we can rewrite Eq. (9) in Kirkpatrick’s notation:

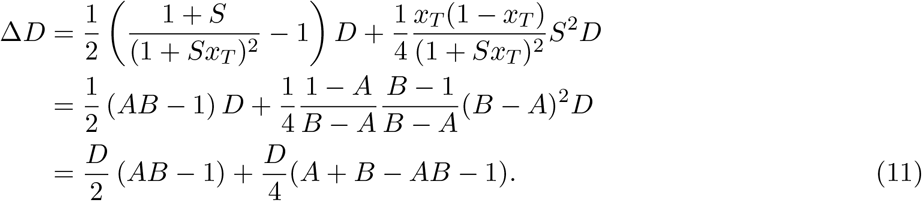

The second term is the contribution of the sexual admixture mechanism. Combining the two terms in Eq. (11) gives

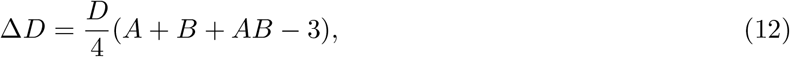

which is Eq. 1c in Kirkpatrick (1982) when *r* = 0.

## Footnotes

1 The calculation for LD in the next generation is 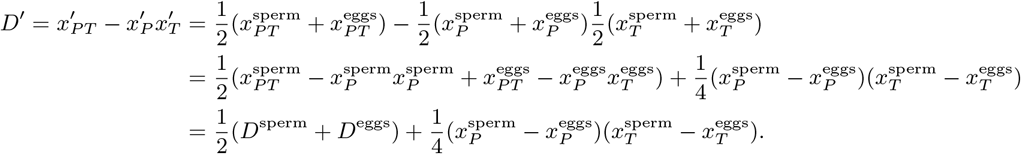

## References

Barton, N. H., Turelli M. (1991). Natural and sexual selection on many loci. Genetics, 127(1), 229–255.

Bay R.A., Arnegard M.E., Conte G.L., Best J., Bedford N.L., McCann S.R., … Schluter D. (2017). Genetic coupling of female mate choice with polygenic ecological divergence facilitates stickleback speciation. Current Biology, 27(21), 3344–3349.

Chenoweth S.F., Blows M.W. (2006). Dissecting the complex genetic basis of mate choice. Nature Reviews Genetics, 7(9), 681.

Crow J.F. (1988). The importance of recombination, pp 56–73. In R. E. Michod and B. R. Levin (eds.), The Evolution of Sex: An Examination of Current Ideas. Sinauer, Sunderland.

Curtsinger J.W., Heisler I.L. (1988). A diploid “sexy son” model. American Naturalist, 132(3), 437–453. Dawkins R. (1976). The Selfish Gene. Oxford University Press, Oxford

Dawkins R. (1986). The Blind Watchmaker. W. W. Norton & Company, New York

Edward D.A., Chapman T. (2011). The evolution and significance of male mate choice. Trends in Ecology & Evolution, 26(12), 647–654.

Faria G.S., Varela S.A., Gardner A. (2018). The relation between RA Fisher’s sexy-son hypothesis and WD Hamilton’s greenbeard effect. Evolution Letters, 2(3), 190–200.

Fisher R.A. (1930). The Genetical Theory of Natural Selection. The Clarendon Press, Oxford

Fitzpatrick C.L., Servedio M.R. (2018). The evolution of male mate choice and female ornamentation: a review of mathematical models. Current Zoology, 64(3), 323–333.

Gomulkiewicz R.S., Hastings A. (1990). Ploidy and evolution by sexual selection: a comparison of haploid and diploid female choice models near fixation equilibria. Evolution, 44(4), 757–770.

Greenspoon P.B., Otto S.P. (2009). Evolution by Fisherian sexual selection in diploids. Evolution, 63(4), 1076–1083.

Heisler I.L., Curtsinger J.W. (1990). Dynamics of sexual selection in diploid populations. Evolution, 44(5), 1164–1176.

Kirkpatrick M. (1982). Sexual selection and the evolution of female choice. Evolution, 36(1), 1–12.

Kirkpatrick M. (1985). Evolution of female choice and male parental investment in polygynous species: the demise of the “sexy son”. American Naturalist, 125(6), 788–810.

Kirkpatrick M. (1987). The evolutionary forces acting on female mating preferences in polygynous animals, pp 67–82. In J. W. Bradbury and M. B. Andersson (eds.), Sexual Selection: Testing the Alternatives. John Wiley and Sons, Chichester.

Kirkpatrick M., & Barton N. (2006). Chromosome inversions, local adaptation and speciation. Genetics, 173(1), 419–434.

Kirkpatrick M., Johnson T., Barton N. (2002). General models of multilocus evolution. Genetics, 161(4), 1727–1750.

Kronforst M.R., Young L.G., Kapan D.D., McNeely C., O’Neill, R. J., Gilbert L.E. (2006). Linkage of butterfly mate preference and wing color preference cue at the genomic location of *wingless*. Proceedings of the National Academy of Sciences, 103(17), 6575–6580.

Kuijper B., Pen I., Weissing F.J. (2012). A guide to sexual selection theory. Annual Review of Ecology, Evolution, and Systematics, 43, 287–311.

Lande R. (1981). Models of speciation by sexual selection on polygenic traits. Proceedings of the National Academy of Sciences, 78(6), 3721–3725.

Mable B.K., Otto S.P. (1998). The evolution of life cycles with haploid and diploid phases. BioEssays, 20(6), 453–462.

Muralidhar P. (2019). Mating preferences of selfish sex chromosomes. Nature, 570: 376–379.

Otto S.P. (1991). On evolution under sexual and viability selection: a two-locus diploid model. Evolution, 45(6), 1443–1457.

Pizzari T., Gardner A. (2012). The sociobiology of sex: inclusive fitness consequences of inter-sexual interactions. Philosophical Transactions of the Royal Society B: Biological Sciences, 367(1600), 2314–2323.

Pomiankowski A. (1987). The costs of choice in sexual selection. Journal of Theoretical Biology, 128(2), 195–218.

Pomiankowski A., Iwasa Y., Nee S. (1991). The evolution of costly mate preferences I. Fisher and biased mutation. Evolution, 45(6), 1422–1430.

Seger J. (1985). Unifying genetic models for the evolution of female choice. Evolution, 39(6), 1185–1193.

Servedio M., Bürger R. (2018). The effects on parapatric divergence of linkage between preference and trait loci versus pleiotropy. Genes, 9(4), 217.

Shaw K.L., Lesnick S.C. (2009). Genomic linkage of male song and female acoustic preference QTL underlying a rapid species radiation. Proceedings of the National Academy of Sciences, 106(24), 9737–9742.

Úbeda F., Haig D., Patten M.M. (2010). Stable linkage disequilibrium owing to sexual antagonism. Proceedings of the Royal Society B: Biological Sciences, 278(1707), 855–862.

Veller C., Kleckner N., Nowak M.A. (2019). A rigorous measure of genome-wide genetic shuffling that takes into account crossover positions and Mendel’s second law. Proceedings of the National Academy of Sciences, 116(5), 1659–1668.

Xu M., Shaw K.L. (2019). Genetic coupling of signal and preference facilitates sexual isolation during rapid speciation. bioRxiv. doi: https://doi.org/10.1101/694497

